# Excess False Positive Rates in Methods for Differential Gene Expression Analysis using RNA-Seq Data

**DOI:** 10.1101/020784

**Authors:** David M. Rocke, Luyao Ruan, Yilun Zhang, J. Jared Gossett, Blythe Durbin-Johnson, Sharon Aviran

## Abstract

**Motivation:** An important property of a valid method for testing for differential expression is that the false positive rate should at least roughly correspond to the p-value cutoff, so that if 10,000 genes are tested at a p-value cutoff of 10^−4^, and if all the null hypotheses are true, then there should be only about 1 gene declared to be significantly differentially expressed. We tested this by resampling from existing RNA-Seq data sets and also by matched negative binomial simulations.

**Results:** Methods we examined, which rely strongly on a negative binomial model, such as edgeR, DESeq, and DESeq2, show large numbers of false positives in both the resampled real-data case and in the simulated negative binomial case. This also occurs with a negative binomial generalized linear model function in R. Methods that use only the variance function, such as limma-voom, do not show excessive false positives, as is also the case with a variance stabilizing transformation followed by linear model analysis with limma. The excess false positives are likely caused by apparently small biases in estimation of negative binomial dispersion and, perhaps surprisingly, occur mostly when the mean and/or the dispersion is high, rather than for low-count genes.

**Supplementary Information:** The computational tools developed for this study are freely available via our website http://dmrocke.ucdavis.edu/software.html. They can be downloaded as R code or run directly through an interactive web-based shiny application to reproduce the analysis presented here per a user’s choice of dataset and the methods to be evaluated.

## 1 INTRODUCTION AND BACKGROUND

### 1.1 Gene Expression

Transcriptome profiling is a powerful means of characterizing a cell’s state and as such, is key to elucidating cellular processes, dissecting genotype-phenotype relationships, and understanding health and disease. As genes are expressed at levels that can vary greatly both within a cellular sample and between samples, it is imperative not only to quantify these expression levels robustly but also to detect and assess between-sample differences with high fidelity.

For nearly two decades, cDNA microarrays have been the platform of choice for profiling complex mixtures of RNAs sampled from the transcriptome (Bertone et al. (2004), Subramanian et al. (2005)). Microarray technology transformed and accelerated gene expression studies by enabling large-scale and genome-wide measurements via parallel, automated, and cost-efficient hybridization-based detection of nucleic acids. While it accurately quantifies relative prevalence of transcripts that derive from a set of preidentified genes, this technology also suffers major limitations. Primarily, it cannot support exploratory studies or facilitate *de novo* discoveries, as it requires *a priori* design and fabrication of the array with the genes that are to be probed. Such considerations essentially preclude unannotated genes or unknown splice variants from being detected as well as limit the number of variants that a single experiment can assay. Other issues include limited dynamic range and imperfections in signal acquisition, which warrant dedicated and non-trivial analysis methodology.

A fundamentally new approach to large-scale and comprehensive RNA analysis has emerged with the recent advent of nextgeneration DNA sequencers (Wold and Myers (2008)). While this technological advance was initially directed at massive genome sequencing, it was soon harnessed to provide sequencing-based RNA transcript quantification, dubbed RNA-Seq (Mortazavi, Williams, McCue, Schaeffer, and Wold (2008), Nagalakshmi et al. (2008)). RNA-Seq was rapidly adopted and standardized due to the assay’s simplicity, affordability, enhanced information content, and scalability; it has become standard practice in molecular biology and biomedical research (Van Keuren-Jensen, Keats, and Craig (2014) S. Li et al. (2014)). This paradigm shift clearly contributed to the scaling trajectory of gene expression studies, but most importantly, it expanded their scope by allowing detection of transcripts *de novo* and at very low abundances. Yet, such unprecedented level of detail also presents multiple new informatics challenges, in particular with respect to transcript reconstruction, quantification, and comparative or differential analysis. In this paper, we show that certain methods used to test for differential expression in RNA-Seq have very poor type I error properties.

### 1.2 Statistical Tests for Differential Expression

Perhaps the most fundamental choice in a pipeline for differential expression analysis is the basic statistical test used for each gene. Many packages and methods, including DESeq ((Anders & Huber, 2010); Anders and Huber (2012)), DESeq2 (Love, Huber, and Anders (2014)), edgeR (Robinson, McCarthy, and Smyth (2010), McCarthy, Chen, and Smyth (2012), Robinson and Smyth (2007), Robinson and Smyth (2008)) and edgeR-robust (X. Zhou, Lindsay, and Robinson (2014)), are based on a negative binomial model for the counts in the expression matrix and explicitly use the negative binomial likelihood. The mathematics of this is detailed in most of the papers supporting these methods, but the rationale seems to be as follows:

1. RNA-Seq data are in the form of counts.
2. The simplest model for count data is the Poisson, in which the mean and the variance are the same.
3. But in most cases, the data are over-dispersed, meaning that the variance is greater than the mean. This may occur when individual observations for a given gene under given conditions are each Poisson, but with a mean that varies from observation to observation.
4. Perhaps the simplest distribution for over-dispersed count data is the negative binomial, which is obtained when each observation is Poisson but the mean varies from sample to sample following a gamma distribution.

One consequence of this assumption is a relationship between the mean and the variance. If a random variable X has a negative binomial distribution with mean μ and variance σ^2^, then σ^2^ = μ + θμ^2^, where θ can be called the dispersion parameter.

The limma package (Smyth (2005)), originally developed for gene expression microarray analysis, is based on fitting a standard linear model for each gene. By default, in a standard linear model, the error variance is the same for all observations from the same gene, for treatment or control. After a log-like transformation (B. P. Durbin, Hardin, Hawkins, and Rocke (2002); B. Durbin and Rocke (2003);B. P. Durbin and Rocke (2004); Rocke and Durbin (2001); Rocke and Durbin (2003)), this might be a plausible assumption for microarray data, though care in the choice of the transformation is important. For RNA-Seq data also, the variance tends to rise with the mean in a very marked fashion, so that the assumption of equal variance cannot hold in general without a transformation.

Taking logarithms can help with this problem, but it fails for low counts, especially zero counts. For the basic treatment vs. control comparison, limma conducts what is actually a pooled two-sample t-test, which assumes equal variance. In order to handle the problem of variances trending with the mean, the voom function (Law, Chen, Shi, and Smyth (2014)) was written as a front end for limma, yielding a method called limma-voom. The voom function calculates variance-based weights that, together with a started log transformation, are meant to stabilize the variance.

In addition, we consider a straightforward negative-binomial-based statistical test arising from the generalized linear model regression formulation with the negative binomial distribution as implemented in the R function glm.nb() in the MASS package (Ripley (2011)). This is the likelihood ratio test (Neyman and Pearson (1928)) which compares the model to a simpler model, which is similar to the ANOVA F-test in ordinary linear regression (Snedecor (1934)). In the case of a two-condition experiment, we compare a model with the condition as a predictive factor to one that has only the mean as a predictor. We could also use the Wald test on the coefficient, though the performance of this test is even worse than that of the likelihood ratio test.

We will also consider the application of a transformation to the counts before applying a test. As described in Bartlett’s 1947 paper (Bartlett (1947)), once a variance function (relationship between the variance and the mean) has been defined, one can derive a transformation that approximately stabilizes the variance. After some derivation it can be shown that if σ^2^ = μ + θμ^2^, then the transformation

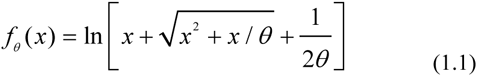

approximately stabilizes the variance, meaning that the variance no longer depends systematically on the mean. Thus, after transformation, we can use methods that don’t explicitly model the variance as a function of the mean, such as linear regression, the analysis of variance, or in the simple two-condition case, the pooled ttest. One way to estimate the best value of θ is to regress the geneand-condition-specific variance on the gene-and-condition-specific mean, and find a value of θ for which the variance neither increases systematically with the mean nor decreases systematically. We can operationalize this by finding a value of θ for which the slope of the regression line is zero. This procedure can be used instead of the voom function and then the data can be submitted to limma. We refer to this procedure as limma-trans.

## 2 METHODS—MEASURING PERFORMANCE WHEN THERE ARE NO TRUE GROUP DIFFERENCES

### 2.1 Packages and Estimators Used

This paper is not meant to be a comprehensive review of packages and methods for differential expression analysis of RNA-Seq data. It is instead meant to make a specific point, which is that methods based on estimates of the parameters of the negative binomial distribution can result in highly inflated false positive levels. We chose one commonly used package based on the negative binomial (edgeR), one which does not use that assumption except in the variance function weighting (limma-voom), a positive control (glm.nb) and a negative control (limma-trans). We have run these procedures on other packages and methods such as DESeq2 and edgeR-robust with essentially similar results. Of course, many other methods exist, including but not limited to: Cuffdiff (Trapnell et al. (2010)), Cuffdiff2 (Trapnell et al. (2013)), NBPSeq (Di, Schafer, Cumbie, and Chang (2011)), TSPM (Auer and Doerge (2011)), baySeq (Hardcastle and Kelly (2010)), EBSeq (Leng et al. (2013)), NOISeq (Tarazona, García-Alcalde, Dopazo, Ferrer, and Conesa (2011)), SAMseq (J. Li and Tibshirani (2013)), ShrinkSeq (Van De Wiel et al. (2012)), DEGSeq (Wang, Feng, Wang, Wang, and Zhang (2010)), BBSeq (Y.-H. Zhou, Xia, and Wright (2011)), FDM (Singh et al. (2011)), RSEM (B. Li and Dewey (2011)), Myrna (Langmead, Hansen, and Leek (2010)), PANDORA (Moulos and Hatzis (2014)), ALDEx2 (Fernandes et al. (2014)), PoissonSeq (J. Li, Witten, Johnstone, and Tibshirani (2011)), and GPSeq (Srivastava and Chen (2010)). We provide code that can be easily adapted to any method that runs in R and applied to the publicly available data sets we used, as well as others.

### 2.2 Resampling from Existing Data Sets

We use three publicly available data sets, the Bottomly data (Bottomly et al. (2011)), the Cheung data (Cheung et al. (2010)), and the Pickrell/Montgomery (“MontPick”) data (Pickrell et al. (2010), Montgomery et al. (2010)), as obtained from the recount site (http://bowtie-bio.sourceforge.net/recount/). The Bottomly data consist of RNA-Seq results from 21 samples of gene expression in the striatum in two strains of mice, 10 from the C7BL/6J strain and 11 from the DBA/2J strain. The Cheung data consist of RNA-Seq results from 41 human samples of B-cell gene expression from the HapMap project. These were from the CEPH (Utah residents with ancestry from northern and western Europe) (CEU) subsample and 17 of the samples were female and 24 were male. The MontPick data consist of RNA-Seq results from 129 human samples of B-cell gene expression from the HapMap project. There were 60 from the above-referenced CEU subset and 69 from the Yoruba people in Ibadan, Nigeria (YRI).

Our resampling method is to choose a sample size *n* = *n*1 = *n*2 such that *n*1 + *n*2 is no larger than the smaller of the two group sizes. We then select *n*1 samples from group 1 at random and compare them with *n*2 samples randomly selected from the same group. Since there are no systematic differences, we should find that the fraction significant is within chance range of the nominal significance level. With, say, 10,000 genes, we should have on the order of 1 gene significant at p = 10^−4^. Of course we could be unlucky and the 3 vs. 3 could have some factor that is more similar within these random groups than across them, but if we repeat the random selection a sufficient number of times, that should average out. Thus, if these methods are performing as advertised, when we conduct a “null” experiment like this, we should not have substantially more significant genes than is implied by the significance level. In this case, we did this 100 times in group 1, 100 times in group 2, and aggregated the results. For each of the three data sets, we conducted this experiment with *n* = 3, 4, or 5 in group 1 and then also for group 2. For the Cheung data set, we also add an 8 vs. 8 comparison and for the MontPick data set a 10 vs. 10 comparison, since the larger sample sizes allow this. In this case, we sampled each trial independently so there can be duplicate samples in some cases (that is the comparison of *n*1 vs. *n*2 might be duplicated. We used this method because it is simple, easy to understand and interpret, and easy to reproduce. If desired, the duplicates could be eliminated with hashing or using the R package partitions (Hankin (2006)).

We then compared the number of nominally significant genes with the expected number of significant genes under the null hypothesis. A method with well-calibrated p-values should have a number of significant genes close to the nominal number. With 10,000 genes and 100 replications in each of the two groups, we have 2 million p-values. The expected number less than 10^−4^ (for example) is then 200.

We chose the 10^−4^ and 10^−6^ levels to make allowance for multiple comparisons. For comparison, with 10,000 genes, the cutoff with the Bonferroni method for a 5% test is 5×10^−6^. We did not use the FDR-adjusted p-values because each of those is a complex function of the entire vector of p-values, and this can make interpretation more difficult, but the raw p-values for a typical method in this data set for FDR-adjusted p-values of 5% are in the range of 10^−4^ and lower. Note that if the null hypothesis is always true, then all genes identified as significant are false positives. If we ask for a set of genes with FDR control at 5%, that means that the number of false positives should be no more than 5% of those identified as significant, and since all genes identified as significant are false positives, that means that an FDR-controlling method should never identify any genes as positive, at least on the average.

### 2.3 Negative Binomial Simulations

In this case, we took one relatively homogeneous data set (the 33 females from the CEU samples in the MontPick data set) and made a crude estimate of the mean μ and dispersion θ of each gene. We estimated μ by the sample mean. If the sample variance was smaller than the mean, we used the Poisson distribution for that gene. Otherwise we estimated θ by

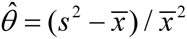

Although this may not be the optimal method for estimating negative binomial parameters, it does not in this case matter, since the only real purpose was to obtain a distribution of means and dispersions that somewhat resembled real data. Before generating the subset, we filtered the MontPick data to eliminate any row with a mean count of less than 1 or where the variance of the CEU female counts were 0, leaving 8122 genes. Clearly, the data set used to obtain the means and variances of the counts is not highly important. We did run the procedure on the other three subsets of the MontPick data with similar results.

We then repeatedly generated negative binomial data (or very occasionally Poisson) resulting in repeated data sets of size 8122 by 33. We did this 100 times.

## 3 RESULTS

### 3.1 Results for resampling from existing data sets

Figure 1 shows the results for the resampling experiment with the MontPick data set and n = 5. The dramatic departures from the null can be seen for edgeR and the likelihood ratio test from the R function glm.nb. As mentioned previously, the Wald test from glm.nb is even worse.

**Fig. 1.**
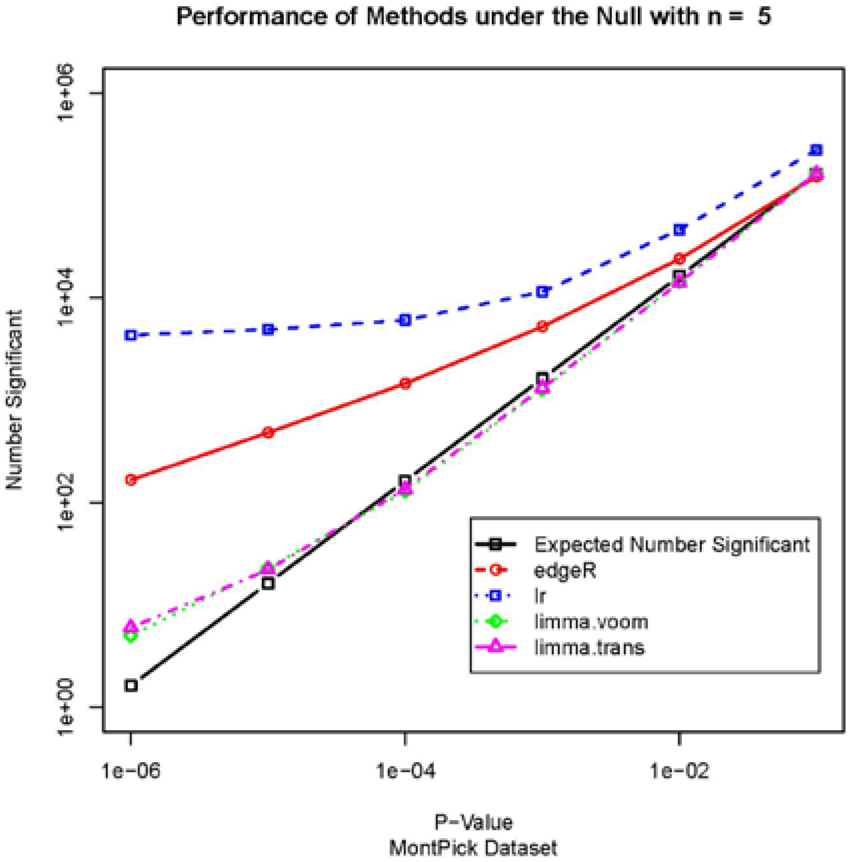
False positives by four methods for a null simulation with the MontPick data set.

Tables 1 and 2 show the actual number of false positives at p = 10^−4^ and p = 10^−6^ for resampling from the four estimators. It can be seen that edgeR and the likelihood ratio test with glm.nb have very large inflation of false positives. There are a number of possible explanations for this, but, as we show next, it is unlikely that these discrepancies arise from special properties of the data sets, or from our resampling approach.

**Table 1.** False positives at p = 10^−4^ for four estimators in null simulations obtained by resampling three data sets.

| Data Set | n | Expected | edgeR | Limma<br>voom | Limma<br>trans | lr |
| --- | --- | --- | --- | --- | --- | --- |
| Bottomly | 3 | 237 | 4560 | 1110 | 508 | 2790 |
| Bottomly | 4 | 237 | 1103 | 124 | 200 | 951 |
| Bottomly | 5 | 237 | 2155 | 571 | 426 | 1328 |
| Cheung | 3 | 181 | 3798 | 198 | 326 | 14113 |
| Cheung | 4 | 181 | 2634 | 123 | 171 | 9006 |
| Cheung | 5 | 181 | 2845 | 200 | 227 | 6359 |
| Cheung | 8 | 181 | 1824 | 117 | 146 | 2707 |
| MontPick | 3 | 162 | 2099 | 144 | 194 | 11537 |
| MontPick | 4 | 162 | 1755 | 168 | 164 | 8953 |
| MontPick | 5 | 162 | 1450 | 129 | 135 | 6047 |
| MontPick | 10 | 162 | 1146 | 139 | 116 | 1752 |

**Table 2.** False positives at p = 10^−6^ for four estimators in null simulations obtained by resampling three data sets.

| Data Set | n | Expected | edgeR | Limma<br>voom | Limma<br>trans | lr |
| --- | --- | --- | --- | --- | --- | --- |
| Bottomly | 3 | 2.37 | 1071 | 41 | 11 | 457 |
| Bottomly | 4 | 2.37 | 160 | 1 | 1 | 391 |
| Bottomly | 5 | 2.37 | 350 | 18 | 4 | 322 |
| Cheung | 3 | 1.81 | 674 | 2 | 12 | 9398 |
| Cheung | 4 | 1.81 | 351 | 0 | 1 | 6350 |
| Cheung | 5 | 1.81 | 353 | 1 | 5 | 4509 |
| Cheung | 8 | 1.81 | 136 | 0 | 0 | 1896 |
| MontPick | 3 | 1.62 | 410 | 5 | 9 | 7045 |
| MontPick | 4 | 1.62 | 307 | 5 | 7 | 6001 |
| MontPick | 5 | 1.62 | 166 | 5 | 6 | 4319 |
| MontPick | 10 | 1.62 | 75 | 1 | 0 | 1215 |

### 3.2 Results from Negative Binomial Simulations

Table 3 shows the number of false positives at p = 10^−4^ in 1000 simulated null negative binomial data sets, each with 8122 genes for four sample sizes and for edgeR and limma-voom. It is apparent that the number of false positives for edgeR is large even under the negative binomial distribution, though the number does decline as *n* gets larger. Limma-voom (and limma-trans, not shown) do not have large numbers of false positives.

**Table 3.** False positives at p = 10^−4^ for edgeR and limma-voom in negative binomial simulated data.

| n | Expected | edgeR | limma-voom |
| --- | --- | --- | --- |
| 3 | 812 | 7138 | 541 |
| 4 | 812 | 6311 | 949 |
| 5 | 812 | 5847 | 1062 |
| 10 | 812 | 3575 | 915 |

An interesting characteristic of p-values from edgeR is that the false-positive p-values are not just small, many of them are extremely small. A histogram of the p-values (say for *n* = 5) does not show any apparent excess of false positives (not shown). Only 8% of the p-values are less than 0.10 instead of an expected 10%. But if we examine the p-values that are less than 0.10, then 15% of them are less than 0.01 (with again an expected fraction of 10%). This continues all the way down. Of the p-values less than 10^−7^, 54% are less than 10^−8^. Thus, there are a large number of very small p-values.

### 3.3 Which Genes are Generating the False Positives?

Under the negative binomial, the only parameters determining the distribution of counts are the mean μ and dispersion θ. It is commonly assumed that it is the low-count genes that cause trouble in the analysis, and it is true that there are often convergence problems in genes with low mean counts, but the false positives are arising from the other end of the parameter distribution.

In Table 4, we have partitioned the means in the CEU female data set into quintiles, and also partitioned the dispersions in to quintiles. The row headings show the geometric mean of the sample means in the quintile and the column headings show the square root of the mean of the estimated dispersions in the quintile. The number in each cell is the number of false positives at p = 10^−4^ for genes in that cell divided by the number of genes falling into that cell for *n* = 5 in 1000 trials of edgeR with negative binomial simulated data. Under the null, the numbers in each cell should be approximately 0.1—the shaded cells are the ones more than twice the nominal.

**Table 4.** False positive ratios at p = 10^−4^ for edgeR in negative binomial simulated data partitioned by quintiles of the mean and dispersion. The row headings show the geometric mean of the sample means in the quintile and the column headings show the square root of the mean of the estimated dispersions in the quintile. The number in each cell is the number of false positives at p = 10^−4^ for genes in that cell divided by the number of genes falling into that cell for n = 5 in 1000 trials of edgeR with negative binomial simulated data. Under the null, the numbers in each cell should be approximately 0.1—the shaded cells are the ones more than twice the nominal.

| Mean | Dispersion |  |  |  |  |
| --- | --- | --- | --- | --- | --- |
|  | 0.4 | 0.5 | 0.6 | 0.9 | 1.7 |
| 1 | 0 | 0 | 0 | 0.011 | 0.198 |
| 8 | 0 | 0 | 0.024 | 0.154 | 3.24 |
| 32 | 0 | 0.015 | 0.098 | 0.863 | 10.676 |
| 108 | 0.014 | 0.070 | 0.358 | 2.370 | 11.848 |
| 489 | 0.032 | 0.267 | 0.926 | 3.583 | 6.909 |

What we can see from this table is that the false positives are entirely generated from genes with high means, high dispersions, or both. An implication of this is that the p-value distribution under then null is quite different for genes with different means and dispersions, so p-values obtained by edgeR and other negativebinomial-based estimators cannot be used to rank genes. Consider the 147 genes that fall into the second quintile in mean and the second quintile in dispersion. Out of 147,000 statistical tests, none is significant at the p = 10^−4^ level, instead of an expected 15. On the other hand, the 146 genes that fall into the fourth quintile in the means and the fourth quintile in the dispersions had 346 tests significant at the p = 10^−4^ level. This means that genes in the higher quintiles of means and dispersions are far more likely to be top genes than those in the lower quintiles, all other things being equal, and therefore that the ranking of genes by p-value is unreliable, not just the level of the test.

## 4 DISCUSSION

There have been numerous reviews and evaluations of methods for RNA-Seq data, many quite comprehensive, and a number of them have compared the power and type I error rates, with varying conclusions. Soneson and Delorenzi (2013), Kvam, Liu, and Si (2012), and van de Wiel, Neerincx, Buffart, Sie, and Verheul (2014) compared the null performance of several methods and found poor control of false discovery rate (FDR) in general. Robles et al. (2012) likewise compared 3 methods and found inflated type I error rates for some methods and conservative performance from others. Reeb and Steibel (2013) compared 3 methods using “plasmodes” (resampled data) and found inflated type I error rates for small significance levels. Guo, Li, Ye, and Shyr (2013) compare 6 methods and conclude that all “suffer from over-sensitivity”. However, Rapaport et al. (2013) compared 7 different methods and concluded that most methods provide good FDR control at the 5% level. Finally, Nookaew et al. (2012) compared 5 different methods with each other and with microarray data and concluded that agreement was good between methods and platforms, and Guo et al. (2013) concluded that agreement between the methods compared was “reasonable”.

Mostly, the examination of Type I error control has been limited to p-values such as 0.05 which are important in analysis of a single outcome. But when testing 10,000 genes, Type I error for small p-values such as 10^−4^ or 10^−6^ are more important. Control of the false discovery rate after multiplicity adjustment has also been frequently examined, but this is harder to interpret since each individual FDR-adjusted p-value (q-value) depends on the entire vector of p-values. In addition, if all the null hypotheses are true, then the actual false discovery rate of any non-empty set of “significant” genes is 100%.

Most likely, the phenomenon of excess false positives in methods based strongly on the negative binomial distribution is a consequence of the known downward bias in the MLE for the negative binomial dispersion (Piegorsch, 1990). While this bias is generally small when viewed from the point of view of parameter estimation, it apparently has a large effect on the tail areas of the test statistic. In any case, it appears that the degree of violation of Type I error control in many existing methods has been greatly underestimated. For the time being, it seems safer to use methods like limma-voom that do not depend explicitly on the negative binomial likelihood.

## AUTHORS’ CONTRIBUTIONS

DMR directed the study, did some of the computations, and was primary author of the manuscript. LR wrote much of the code and performed most of the simulation experiments. JJG and YZ each wrote some of the code, and did bibliographic research. BDJ and SA wrote parts of the manuscript and edited it and did background research. All participants contributed to the research plan and the outline of the manuscript in extensive and frequent meetings and discussions.

## FUNDING

This work was supported by National Institutes of Health (NIH) grants R33AI080604 and UL1 RR024146 (David Rocke) and R00 HG006860 (Sharon Aviran)

